# High throuput label free rapid screening to identify cell penetrating peptide

**DOI:** 10.1101/2025.05.19.654248

**Authors:** Vivek Kumar

## Abstract

Fluorophore-peptide tagging is a widely used method for tracking peptide. However, this approach is slow and costly, which hinders the design and screening of native cell penetrating peptide. The extrinsic fluorophore ANS (1-anilino-8-naphthalene sulfonate) can monitor change in membranes hydrophobicity during the internalization of cationic peptide. This property makes ANS a potentially attractive low cost tool for high throughput rapid screening of native peptides with cellular penetration property.

## Introduction

Peptides have emerged as promising biomolecules for drug/macromolecules delivery and therapeutic applications due to their biocompatibility and specificity ^1^. A crucial aspect of peptide-based therapeutics is understanding their cellular penetration properties, which are often studied using fluorescence labelling techniques. Traditionally, this requires covalent attachment of fluorophores to peptides, a process that can incur huge cost of fluorophore tagging and technical expertise and significant time. Additionally, tagging with a fluorophore can slow down the screening process, limiting the number of peptides that can be screened.

Here, we present a method that enables rapid screening of native peptides facilitating the use of unmodified peptides in cellular studies. We used the extrinsic fluorophore, 1-anilino-8-naphthalene sulfonate (ANS) to screen the cell-penetrating ability of several peptide sequences. ANS is a fluorescent probe that binds to the hydrophobic region of proteins. It does not fluoresce in water, but there is a shift of emission maxima from 545 nm for free ANS to 470 nm for its bound form, that further increases after the formation of the protein-ANS complex^2–5^ANS interacts with the anion permeable region of the membrane^6^, and this binding of ANS to the plasma membrane leads to a blue shift in the peak, causing a 4-fold increase in fluorescence intensity^7^. It is well known that there is little change in fluorescence with time, and it appears as soon as the ANS is added to the cell^2^. The internalization of well-established cationic cell-penetrating peptides R9 and (TAT-_49-57_) through the apolar (hydrophobic) region of plasma membrane of wheat protoplasts and animal cells (HEK 293T) was used as positive control, while a known non-cell-penetrating peptide mTAT (AKKRRQRRR) was used as negative control. This study introduces a non-covalent fluorescent screening method for native peptides, eliminating the need for costly fluorophore tagging.

## Materials & Methods

### Protoplast isolation & Assay

Protoplasts were isolated from 7 days old wheat plants following a previously described protocol ^1^. Protoplasts were pre-incubated (10 minutes) with 100 µM ANS and then treated with different peptides R9 (10 µM), TAT (10 µM) and mTAT (5 µM) as per reaction setting. Change in fluorescence were determined by fluorescence plate reader after 10 minutes and 20 minutes. Buffer used was CPW buffer.

### Mammalian Cell Assay (HEK 293T)

HEK 293T cells were cultured as per defined protocol. Briefly, HEK cells were grown in DMEM supplemented with 10%. For assay setting, HEK cells were washed with 1X PBS and then cell count was adjusted to 10^5^ cells/ml. 100 µl of HEK cells were pre-incubated (10 minutes) with 100 µM ANS followed by peptide treatments of R9 (10 µM) and mTAT (5 µM). Buffer used was 1X PBS. Change in fluorescence was determined by fluorescence plate reader after 10 minutes and 20 minutes. Buffer used was 1X PBS.

## Results Protoplast Assay

Treatment with the cationic peptide R9 (10 µM) led to a decrease in ANS fluorescence intensity, indicating internalization of the peptide and potential displacement or quenching of ANS due to membrane interaction and pore formation (Figure 1a). In contrast, treatment with the TAT peptide (10 µM) caused an increase in ANS fluorescence, suggesting enhanced binding of ANS to newly exposed hydrophobic regions, consistent with peptide penetration and membrane disruption (Figure 1b). The non-penetrating control peptide (mTAT-AKKRRQRRR) (5 µM) did not produce any significant change in ANS fluorescence, indicating a lack of membrane interaction or peptide internalization (Figure 1c). Buffer fluorescence and peptide fluorescence shows no background fluorescence (Figure 1d). Demonstrate that ANS fluorescence responds differentially to membrane activity induced by penetrating versus non-penetrating peptides in plant cells.

**Figure 1.**
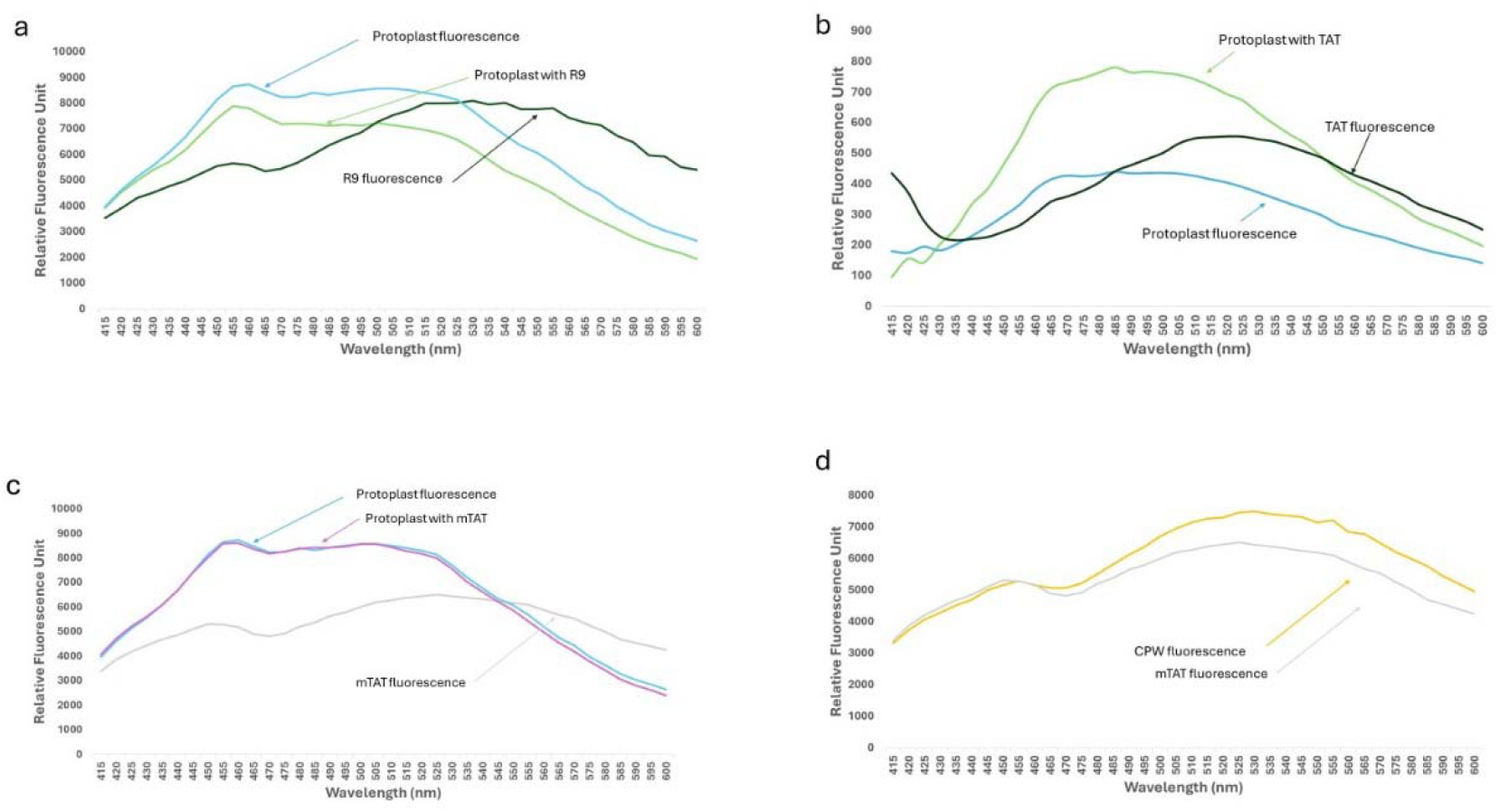
ANS fluorescence on translocation of the cell-penetrating peptide in wheat protoplast. Protoplasts were pre incubated (10 minutes) with ANS treated with (a) R9 (10 µM), (b) TAT (10 µM) and (c) mTAT (5 µM) peptide. Buffer use to incubate wheat protoplast was CPW buffer. ANS concentration used was 100 µm.

### Mammalian cell Assay (HEK 293T)

Treatment with R9 (10 µM) led to a notable increase in ANS fluorescence, which contrasts with its effect in protoplasts. This may reflect differences in membrane composition or peptide interaction dynamics in mammalian cells (Figure 2a). Similar to protoplasts, treatment with mTAT (5 µM) did not alter ANS fluorescence, reaffirming its inability to penetrate cell membranes (Figure 1c & 2b). These findings suggest that ANS-based screening can differentiate peptide internalization behaviour in a cell type-dependent manner, highlighting its utility across diverse biological systems.

**Figure 2.**
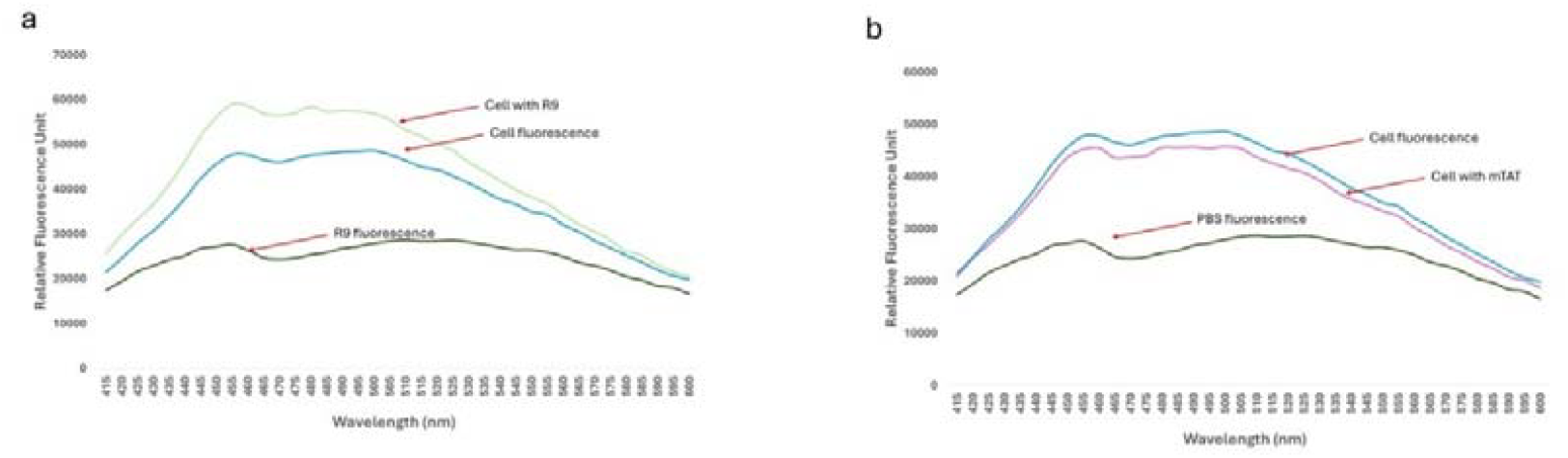
ANS fluorescence on translocation of the cell-penetrating peptide in HEK 293T cells. HEK 293T were pre incubated (10 minutes) with ANS with (a) R9 (10 µM) and (b) mTAT (5 µM). Buffer use to incubate HEK 293T cells was 1X PBS. ANS concentration used was 100 µm.

## Discussion

TAT _(49-57),_ a peptide derived from the HIV-1 TAT protein, and polyarginine (R9) are well-known for their ability to efficiently translocate into both plant protoplasts and animal cells, where they accumulate in the nucleus ^8,9,10^. Treatment with ANS induces a blue shift in its fluorescence peak, indicating changes in the local environment. Further treatment of ANS-pretreated cells with cationic peptides resulted in altered fluorescence intensity, which varied depending on the peptide used and cell type (Fig. 1) and an indicator of membrane interaction and internalization.

The observed changes in ANS fluorescence may result from the neutralization of negative charges on the membrane surface by cationic peptides, leading to transient alterations in membrane hydrophobicity near the peptide-binding region. This process likely facilitates the formation of transient pores 11, which in turn may influence the binding of ANS molecules to deeper hydrophobic regions within the membrane 12 .

Cationic peptide treatment alters membrane hydrophobicity, as evidenced by the blue shift in the ANS fluorescence peak and changes in fluorescence intensity. Once transient pores form, peptides interact with the inner membrane surface and internalize, ceasing further changes in fluorescence due to shifts in the intracellular environment. The enhanced fluorescence observed upon CPP addition is likely due to modifications in membrane properties, increasing the availability of hydrophobic sites for ANS binding during CPP internalization12.

Therefore, ANS could serve as a novel extrinsic fluorescent probe to analyze unprecedented changes in cell membrane hydrophobicity during the internalization of cationic peptides. ANS acts as a binary indicator, signaling either “ON” or “OFF.” “ON” indicates cellular uptake of the peptide through transient pores or membrane bilayer disruption, while “OFF” suggests no cellular internalization or interaction. This approach provides a rapid method for screening native peptides (without covalently attached fluorophores), bringing us closer to utilizing unmodified peptides in cellular studies. The methodology is applicable to both plant and animal cells and works with cationic as well as hydrophobic peptides. Additionally, this study is particularly valid for peptides that lack tryptophan residues. In contrast, peptides containing tryptophan will exhibit both fluorescence and membrane binding, complicating result interpretation.

## Summary

This study presents a novel label-free, high-throughput screening method for identifying cell-penetrating peptides using the extrinsic fluorophore ANS. The technique successfully distinguishes between penetrating and non-penetrating peptides in both plant and mammalian systems. The use of ANS fluorescence changes as a binary readout offers a cost-effective and rapid alternative to conventional fluorophore tagging, facilitating the study and design of native peptides for drug delivery and therapeutic purposes.

## Statements and Declarations

There is nothing to declare.

## References

1. Kumar V, Chugh A. Cell-penetrating peptide for targeted macromolecule delivery into plant chloroplasts. Appl Microbiol Biotechnol. 2022;106(13-16):5249–5259. doi:10.1007/s00253022-12053-3

2. Rubalcava B, Martínez de Muñoz D GC. Interaction of Fluorescent Probes with Membranes: I. Effect of ions on erythrocyte membranes. Biochemistry. 1969;8(7):2742–2747.

3. Wang N, Faber EB, Georg GI. Synthesis and Spectral Properties of 8-Anilinonaphthalene-1-sulfonic Acid (ANS) Derivatives Prepared by Microwave-Assisted Copper(0)-Catalyzed Ullmann Reaction. ACS Omega. 2019;4(19):18472–18477. doi:10.1021/acsomega.9b03002

4. Latypov RF, Liu D, Gunasekaran K, Harvey TS, Razinkov VI, Raibekas AA. Structural and thermodynamic effects of ANS binding to human interleukin-1 receptor antagonist. Protein Science. 2008;17(4):652–663. doi:10.1110/ps.073332408

5. Matulis D, Lovrien R. 1-anilino-8-naphthalene sulfonate anion-protein binding depends primarily on ion pair formation. Biophys J. 1998;74(1):422–429. doi:10.1016/S0006-3495(98)77799-9

6. Fortes PAG, Hoffman JF. The interaction of fluorescent probes with anion permeability pathways of human red cells. J Membr Biol. 1974;16(1):79–100. doi:10.1007/BF01872408

7. Dionisi O, Galeotti T, Terranova T, Arslan P, Azzi A. Interaction of fluorescent probes with plasma membranes from rat liver and morris hepatoma 3924A. FEBS Lett. 1975;49(3):346–349. doi:10.1016/0014-5793(75)80782-4

8. Vivès, Erice, Priscille Brodin BL. A Truncated HIV-1 Tat Protein Basic Domain Rapidly Translocates through the Plasma Membrane and Accumulates in the Cell Nucleus. J Biol Chem. 1997;272(25):16010–16017. doi:10.1074/JBC.272.25.16010

9. Chugh A, Eudes F. Translocation and nuclear accumulation of monomer and dimer of HIV-1 Tat basic domain in triticale mesophyll protoplasts. Biochim Biophys Acta Biomembr. 2007;1768(3):419–426. doi:10.1016/j.bbamem.2006.11.012

10. Numata K, Horii Y, Oikawa K, Miyagi Y, Demura T, Ohtani M. Library screening of cell-penetrating peptide for BY-2 cells, leaves of Arabidopsis, tobacco, tomato, poplar, and rice callus. Sci Rep. 2018;8(1). doi:10.1038/s41598-018-29298-6

11. Palm-Apergi, C., Lorents, A. P K.,, Pooga, M., and Hällbrink M. The membrane repair response masks membrane disturbances caused by cell-penetrating peptide uptake. The FASEB Journal. 2008;23(1):214–223. doi:10.1096/fj.08-110254

12. Hirose H, Takeuchi T, Osakada H, et al. Transient focal membrane deformation induced by arginine-rich peptides leads to their direct penetration into cells. Molecular Therapy. 2012;20(5):984–993. doi:10.1038/mt.2011.313

